# Hyaluronic Acid-Alginate Hydrazone Crosslinked Hydrogels Support the Generation and Maturation of V2a Interneurons

**DOI:** 10.64898/2026.06.01.729352

**Authors:** Alycia N. Galindo, Madison E. McLaren, Alyssa K. Chi, Jenna D. Khachatourian, Jessica C. Butts, Marian H. Hettiaratchi

**Author notes:** These authors contributed equally to this work (Alycia N. Galindo, Madison E. McLaren).

## Abstract

Injury to the central nervous system (CNS) causes inflammation, cell death, and glial scar formation that inhibits tissue repair. Injectable hydrogels modified with extracellular matrix (ECM)-derived peptides can provide biochemical cues to promote neural tissue repair and serve as a vehicle to deliver therapeutics across the blood-spinal/blood-brain barrier in a minimally invasive manner. We developed an injectable hydrazone crosslinked hyaluronic acid-alginate (HA-Alg) hydrogel for neural tissue repair. We fabricated hydrogels with a range of polymer concentrations and evaluated their physicochemical properties to identify formulations that mimic the stiffness and viscoelastic properties of the CNS tissue environment. Hyaluronic acid was further modified with ECM-derived, cell-adhesive peptides (RGD and IKVAV) to enhance neuronal adhesion and viability. To evaluate the therapeutic potential of our hydrogel platform, we embedded mouse embryonic stem cell aggregates and differentiated them toward mature V2a interneurons. These interneurons are critical for relaying motor signals and represent a promising therapeutic cell population for treating spinal cord injuries. We demonstrated successful enrichment for V2a interneurons in HA-Alg hydrogels containing ECM-derived peptides. Interestingly, both our newly described HA-Alg and established crosslinked HA-HA hydrogels containing IKVAV peptides demonstrated significantly increased neurite length in interneuron enriched cultures compared to hydrogels without peptides. This study demonstrates that the addition of ECM-derived peptides is essential to support the neuronal adhesion and viability required for functional CNS tissue repair.

**Graphical Abstract:** 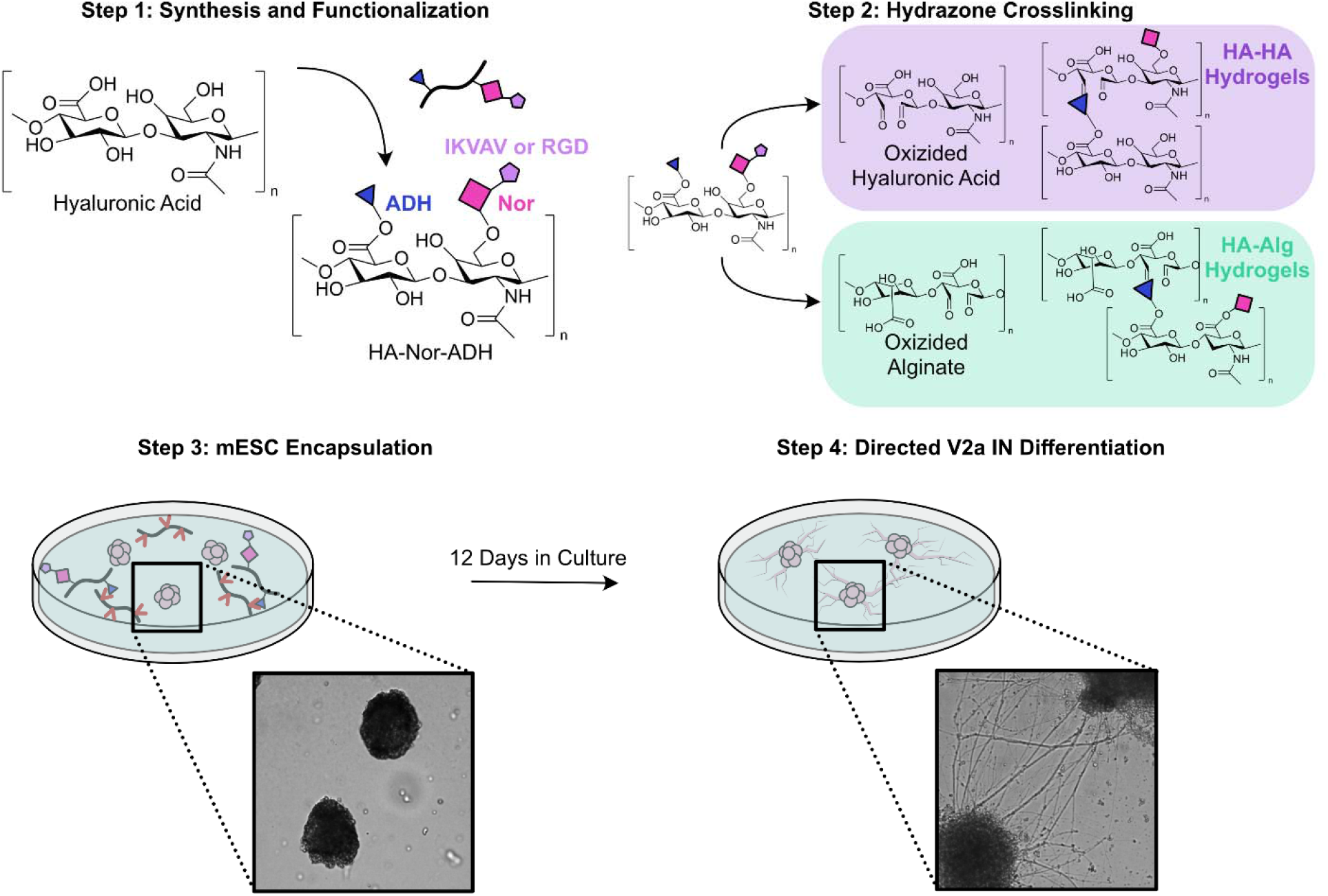

**Statement of Significance:** Development of injectable, peptide-modified hyaluronic acid alginate hydrogels to support neuronal maturation for central nervous system tissue repair.

## Introduction

The spinal cord transmits sensory information to the brain while relaying motor commands back to the peripheral nervous system.^1^ Spinal cord injury (SCI) is characterized by degeneration of neurons that can result in motor, sensory and autonomic dysfunction.^2^ Following injury, the rapid invasion of immune cells and activation of radial glia create an inflammatory environment that causes secondary tissue damage, a cavity of lost tissue, and the formation of a glial scar that physically and biochemically inhibits tissue regeneration.^3^ Various therapeutic approaches have been developed to attempt to restore neural tissue function after SCI. These approaches include targeting the neuroinflammatory environment, reconstituting damaged tissue by delivering cells that support neural tissue repair,^4,5^ removing biomolecules from the injury environment that inhibit tissue repair,^6^ and delivering neurotrophic factors to promote neuronal growth and remyelination.^7,8^ While some of these approaches have had success in promoting neural tissue regeneration in rodent models, these successes rarely translate into meaningful motor or sensory gains in human clinical trials. This translational failure likely stems from the different anatomical scales, the heterogeneity of clinical injuries,^9^ and the development of multi-centimeter cystic cavities that create growth barriers that are magnitudes more severe than those modeled in typical rodent contusions.^9,10^ Biomaterials such as hydrogels, nanoparticles, and aligned scaffolds have the potential to overcome these hurdles, because they can be engineered to support the local delivery of viable cells, sustained release of bioactive proteins, and presentation of other biomolecules that modify the injury environment to support axonal regeneration. Biomaterials can be specifically tailored to treating SCI such that they can be injected through the blood-spinal cord/blood-brain barrier, provide directional topographical cues that guide axons across the lesion, and act as reservoirs that protect therapeutic molecules from rapid degradation *in vivo*.^11^

Hydrogels that mimic mechanical properties of CNS extracellular matrix (ECM), including its stiffness that ranges from 100-1000 Pa^12–15^, cell adhesion, and viscoelastic properties, can support cell proliferation and differentiation and guide axonal extension.^16,17^ Hydrogels made from modified ECM molecules, such as hyaluronic acid (HA), collagen, and gelatin, are particularly attractive biomaterials for treating SCI because they can also mimic the biochemical composition of neural tissue,^18^ providing neuroprotection and stabilizing neuronal networks.^19^ HA, which is the most abundant ECM molecule in the CNS, maintains neural tissue homeostasis by enhancing migration, proliferation, and differentiation of neural progenitor cells and oligodendrocyte progenitor cells to help drive repair after injury.^20,21^ HA can be modified with functional groups such as aldehydes, maleimides, methacrylates, or hydrazides that allow it to be crosslinked into a hydrogel. For these reasons, HA hydrogels have been used extensively in pre-clinical models to deliver cells proteins, and enzymes to the CNS and are a favored biomaterial for CNS tissue engineering applications.^16,22,23^

Alginate, a naturally occurring polysaccharide derived from brown seaweed, is emerging in CNS applications for its antioxidant, anti-inflammatory, and neuroprotective properties.^24,25^ Although not a native component of the brain ECM, alginate hydrogels can be fabricated to match CNS tissue stiffness and viscoelasticity.^26^ Alginate’s high carboxylic acid content allows it to sequester divalent cations, potentially mitigating the oxidative stress triggered by the release of redox-active ions during hemorrhage.^27^ Alginate hydrogels physically crosslinked with calcium ions have facilitated neurite outgrowth and neural stem cell transplantation.^28,29^ However, calcium-crosslinked alginate hydrogels are limited in CNS applications because they can undergo calcium ion exchange, resulting in an inflammatory response,^30,31^ and tend to swell or stiffen in the highly ionic CNS environments.^32^ Alternatively, modifying alginate with functional groups, such as aldehydes, ester linkages, or amide bonds,^33^ can enable chemical crosslinking schemes that do not require the use of calcium ions.

Because functional tissue restoration after SCI depends on the directed regeneration of specific neuronal subpopulations to their natural target regions, we sought to investigate the development of a hydrogel to support the growth and viability of a specific subpopulation of neurons. V2a interneurons are prime candidates for this application, as they are excitatory cells that span the hindbrain and spinal cord, serving as a critical bridge for relaying sensorimotor information between intersegmental regions and contributing to the coordination of sympathetic and parasympathetic functions.^34–36^ Within the spinal cord, these neurons coordinate excitatory inputs that are critical for autonomic and voluntary motor function.^37^ V2a interneurons also play a role in spinal respiratory networks and facilitate the plasticity of neuronal networks in respiratory muscles as observed in models of amyotrophic lateral sclerosis and cervical SCI.^38,39^ The ablation of V2a interneurons in mice disrupts breathing patterns, forelimb reaching tasks, and left-right hindlimb coordination.^36^ The Sakiyama-Elbert lab pioneered initial protocols for V2a interneuron differentiation from mouse embryonic stem cells (mESCs) in 2014.^40,41^ Since then, V2a interneurons have been successfully differentiated from human pluripotent stem cells,^36,37^ and their transplantation in SCI has been identified as a promising therapeutic approach.^42^ V2a interneurons transplanted into cervical SCI rat models functionally integrate with the host spinal cord, improving electrophysical signaling and restoring diaphragm activity.^35,39,43,44^ However, the survival and engraftment of transplanted neurons *in vivo* can be a challenge in larger injuries, as chronic inflammation, the growth-inhibitory environment of the glial scar, and the lack of cell-adhesive substrates can all contribute to cell death.^45^ While HA hydrogels containing astrocyte-derived ECM have been successfully used for the transplantation of V2a interneurons,^44^ neuronal staining showed that neuronal processes in the lesion area was not significantly enhanced in this system compared to HA hydrogels alone, which may be due to the fact that HA inherently lacks neuronal adhesion cues necessary to support neuron attachment and proliferation.^8^ Thus, there is a need to develop hydrogels with cell-adhesive cues to support the viability and engraftment of transplanted neurons.

To address these challenges, here we describe a dual polymer, hyaluronic acid-alginate (HA-Alg) hydrazone crosslinked hydrogel to support V2a interneuron differentiation, maturation and survival for neural tissue repair. We hypothesized that a hydrogel that combines the properties of both HA and alginate through dynamic covalent crosslinking could more closely mimic the biomechanical properties of CNS tissue and eliminate the need for calcium-based crosslinkers. HA was functionalized with adipic acid dihydrazide for crosslinking with oxidized alginate to form reversible hydrazone bonds. This dynamic covalent crosslinking enables the fabrication of injectable hydrogels with shear-thinning and self-healing properties that are ideal for CNS tissue repair as they can be easily injected through the blood-spinal/blood-brain barrier in a minimally invasive manner to fill the cavity of lost tissue.^46–48^ However, despite successfully mimicking the physical properties of CNS tissue, current hydrogels often fail to provide the cell-adhesive cues essential for the long-term survival and maturation of transplanted cells. Thus, we also hypothesized that chemically conjugating ECM-derived peptides, such as fibronectin-derived (RGD) and laminin-derived peptides (IKVAV), to HA could provide an adhesive substrate with the appropriate biochemical cues to support the culture of V2a interneurons.

We evaluated the physicochemical properties of hydrogels fabricated using a range of polymer concentrations to identify a hydrogel formulation that could support neuronal culture. We targeted a hydrogel with a gelation time between 1 and 5 minutes to facilitate cell dispersion throughout the hydrogel, stiffness of ∼100 Pa to mimic CNS tissue, and minimal degradation (±25%) over 28 days to support long-term cell culture. We then investigated whether HA-Alg hydrogels modified with ECM-derived peptides could provide support the differentiation of mESCs into viable, mature V2a interneurons. HA-Alg hydrogels were compared to similar, previously developed HA-HA hydrazone-crosslinked hydrogels to evaluate the use of alginate in a biomaterial for V2a interneuron transplantation. The differentiation and maturity of V2a interneurons were assessed via cell viability, neurite outgrowth and gene expression. We determined that alginate can be combined with HA to form stable hydrogels that meet our criteria for a cell transplantation vehicle and that HA-Alg hydrogels modified with IKVAV can support V2a interneuron viability and maturation. Overall, we developed an HA-Alg hydrogel platform that can be used for future applications in neural tissue repair.

## Materials and Methods

### Synthesis of Norbornene-Hyaluronic Acid (HA-Nor)

HA-Nor was synthesized as previously described.^49,50^ Briefly, an initial intermediate product, tetrabutylammonium-hyaluronic acid (HA-TBA) was created through an ion exchange. HA-TBA was made by dissolving sodium hyaluronic acid (Lifecore Biomedical LLC, Chaska, MN) (1 g, 2.64 mmol) for 20 minutes in 20 mL of double distilled water (ddH_2_O) at 2% w/v. Dowex (Amberlight™ MB) (Sigma Aldrich, Saint Louis, MO) was weighted at a 3:1 mass ratio and added to a flask for ion exchange for 5 hours followed by vacuum filtration. The pH of the filtrate was then adjusted to pH 7 using tetrabutylammonium hydroxide 30-hydrate (Millipore Sigma, Milwaukee, WI). The product was then frozen at -80°C prior to being lyophilized.

HA-TBA (1 g, 1.61 mmol) was dissolved in dimethyl sulfoxide (DMSO) (Fisher Scientific, Waltham, Massachusetts) at 2% w/v for 20 minutes followed by a purge with N_2_ and the addition of 5-norbornene-2-carboxylic acid (Millipore Sigma) at 0.5 molar equivalences (0.11 g, 0.81 mmol), 4-(Dimethylamino)pyridine (DMAP) (Millipore Sigma) Millipore Sigma) at 0.25 molar equivalences (0.05 g,0.40 mmol), and Di-tert-butyl dicarbonate (Boc_2_O) (Millipore Sigma) at 0.07 molar equivalence (0.03 g, 0.11 mmol). The reaction was stirred at 45°C for 20 hours, solution was quenched with equal volume cold water and dialyzed for 3 days against ddH_2_O. Following 3 days, the product was precipitated out with the addition of NaCl (1 g of per 100 mL of solution) and cold acetone (1 L per 100 mL solution). Precipitate was then redissolved in ddH_2_O, frozen, and lyophilized.

### Synthesis of Adipic Acid Dihydrazide Norbornene Hyaluronic Acid (HA-Nor-ADH)

HA-Nor (0.13 g, 0.24 mmol) was then modified a second time with an adipic acid dihydrazide (ADH) group for crosslinking. HA-Nor was dissolved in ddH_2_O at 2% w/v followed by the addition of ADH (Spectrum Chemical Mfg. Corp., Gardena, CA) at 10 molar equivalences (0.418 g, 2.40 mmol) and hydroxybenzotriazole hydrate (Chem-Impex, Wood Dale, IL) at 1 molar equivalence (0.03 g, 0.24 mmol). The pH was adjusted to 4.75, then 1-1-Ethyl-3-[3-dimethylaminopropyl] carbodiimide hydrochloride (EDC) (G-Biosciences, St. Louis, MO) EDC was added at 1 molar equivalence (0.05 g, 0.24 mmol). pH was monitored at 4.75 for 2 hours. The reaction stirred for 48 hours, then dialyzed against 0.1 M NaCl for 2 days followed by dialysis against ddH_2_O for an additional 2 days prior to being frozen and lyophilized.

### Synthesis of Oxidized Alginate (Alg-Ox)

Sodium alginate (low viscosity <20mPa*s and molecular weight <75 kDa) purchased from NovaMatrix (Sandvika, Norway) (1 g, 2.51 mmol) was dissolved at 2% w/v in ddH_2_O. Sodium periodate (NaIO4) (Millipore Sigma) was dissolved in ddH_2_O at 3% w/v at 0.6 molar equivalence (0.32 g, 1.51 mmol) and then added to alginate in a round bottom flask covered with aluminum foil to protect from light. After letting the solution react for 4 hours, the reaction was quenched with 5 molar excess propylene glycol (Avantor, Center, PA) (0.96 g, 12.56 mmol) and rotated for 2 hours. The solution was then dialyzed against ddH_2_O for 2 days, frozen, then lyophilized.^51^ HA was oxidized similarly, as previously described.^50^

### Peptide Conjugation and Quantification

A 1% w/v solution containing HA-Nor-ADH (10 mg, 0.01 mmol, 0.0016 mmol norbornene), 2 molar excess of RGD (CGRGDSG) (GenScript, Piscataway, NJ) (2 mg, 0.0032 mmol) or IKVAV (CGKIKVAVG) (GenScript) (2.8 mg, 0.0032 mmol) to 1 mol of norbornene, 0.05% w/v Irgacure 2959 Advanced Biomatrix (Carlsbad, California), and PBS was made. A thiol-ene reaction between HA-Nor-ADH and the terminal cysteine was performed by exposing the solution to UV light at a wavelength of 365 nm at 5 mW/cm2 for 5 minutes. The solution was dialyzed against ddH2O for 1 day before being sterile-filtered, frozen at -80°C, and lyophilized.

The disappearance of thiols after the photoinitiated reaction was detected using Ellman’s reagent as previously described.^50^ The Ellman’s reagent (Fisher Scientific) consisted of 2 mM of 5,5-dithio-bis-(2-nitrobenzoic acid) (DTNB) and 50 mM sodium acetate in 1 M tris(hydroxymethyl)aminomethane (TRIS) buffer. Samples of the reaction before and after UV exposure were diluted to fall within the range of a standard curve of serially diluted peptides. 50 µL of each standard and sample were added to a 96-well plate and mixed with 250 µL of Ellman’s reagent. The plate was developed at room temperature for 5 minutes prior to reading absorbance at 412 nm on a microplate reader (BioTek Synergy Neo2).

### Degree of Modification (DOM) Determination of HA-Nor and HA-Nor-ADH

Proton nuclear magnetic resonance spectroscopy (H NMR, 500Hz, BrukerUSA) was used to quantify the degree of modification of both HA-Nor and HA-Nor-ADH as previously described.^49,50^ A representative spectrum of HA-Nor-ADH can be found in supplementary information (**Figure S1**).

A hydroxylamine hydrochloride titration was performed to quantify the degree of modification of Alg-Ox. The volume of NaOH used in the titration was used in **equation 1** to equate the degree of modification.

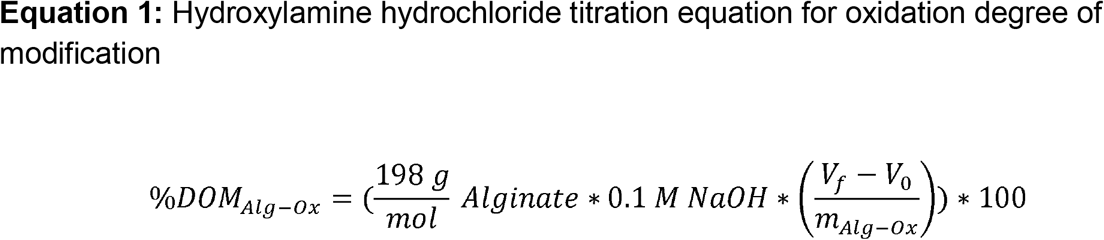

where 198 g/mol is the monomeric molecular weight of alginate, 0.1 M is the molarity of NaOH used to adjust the solution pH, V_f_ is the final volume of NaOH in the burette after titration recorded in liters, V_0_ is the initial volume of NaOH in the burette before titration recorded in liters, and m_Alg-Ox_ is the mass in g of Alg-Ox used in the titration.

### Preparation of Hyaluronic Acid Alginate (HA-Alg) Hydrogels

To form HA-Alg hydrogels to evaluate the gelation time, compressive modulus, and mass change, 50 µL of HA-Nor-ADH and 50 µL Alg-Ox at varying concentrations (1-3% w/v) were mixed **(Table 1)**. Hydrogels used for mass change and gelation time optimization experiments were prepared in 2 mL microcentrifuge tubes. Hydrogels for compression testing were prepared in cylindrical silicon molds with an 8 mm diameter and depth of 4 mm.

**Table 1.**
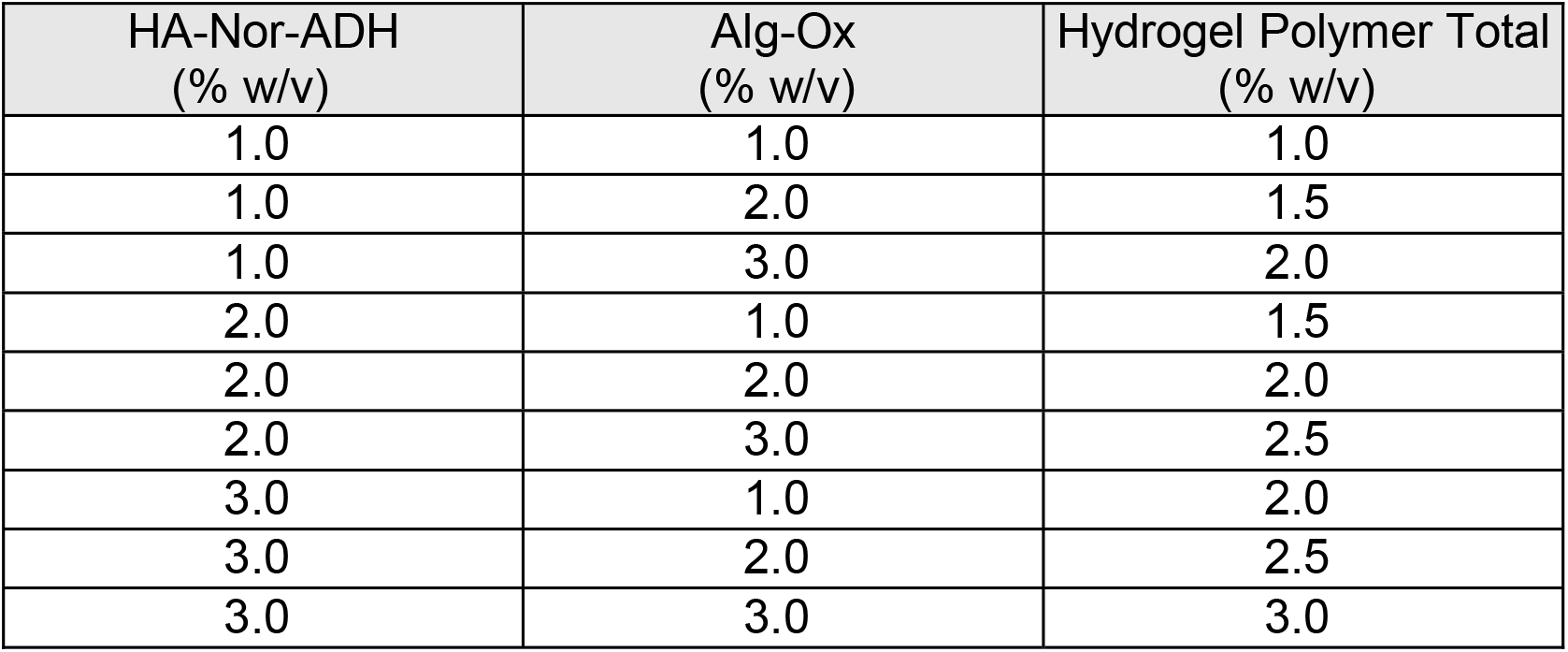
Table of hydrogel formulations for physicochemical characterization.

### Physiochemical Characterization of Hydrogels

#### Gelation Time

The gelation time of the hydrogels were quantified using the inverted tube test.^49^ 50µL of HA-Nor-ADH and 50 µL of Alg-Ox at desired concentrations were added to a 2 mL tube, mixed, and inverted once every minute to assess viscosity until gelation.

Determination of gelation time began at initial mixing of the two solutions and the time there were no visual signs of fluid flow was recorded.

#### Compressive Modulus

Hydrogel compression modulus was evaluated using a Discovery Hybrid Rheometer-2 (DHR-2; TA Instruments, New Castle, DE, USA). 100 µL HA-Alg hydrogels were created in cylindrical molds and placed on a Peltier plate maintained at 37°C. Hydrogels were compressed at a constant linear rate of 10 μm/s to 15% of their original heights using an 8 mm aluminum parallel plate.

#### Mass Change

We determined the mass change of the hydrogels over 28 days in PBS and artificial cerebrospinal fluid (aCSF) (NaCl (149 mM), KCl (3 mM), MgCl_2 *_ 6H_2_O (0.8 mM), CaCl_2 *_ 2H_2_O (1.4 mM), Na_2_HPO_4 *_ 7H_2_O (1.5 mM), and NaH_2_PO_4_ (0.2 mM)). 100 µL hydrogels were formed in 2 mL tubes and allowed to crosslink for 24 hours. Initial crosslinked hydrogels were weighed before the addition of 500µL of 0.05% sodium azide in PBS or 500 µL of aCSF in their respective tubes. At 0 hours, 24 hours, 7, 14, 21, and 28 days, the solution was carefully removed from the tubes, and the hydrogel was weighed. The mass change compared to the initial hydrogel mass was calculated at each time point as a percentage using **equation 2**.

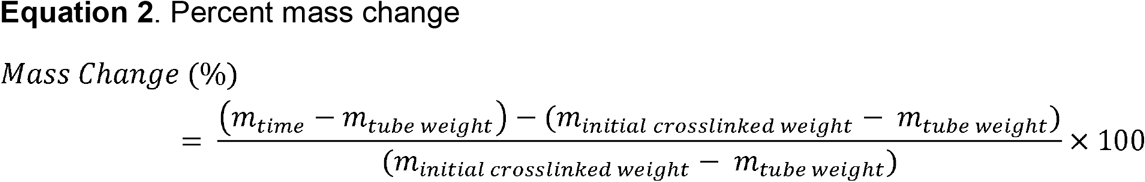

### Ab2.2 Mouse Embryonic Stem Cell Culture

Mouse embryonic stem cells (mESCs) derived from Ab2.2 mice were cultured on gelatin-coated T-25 flasks. The complete culture medium consisted of high-glucose Dulbecco’s Modified Eagle’s Medium (DMEM; Gibco) supplemented with 10% newborn calf serum (NBCS; Gibco), 10% fetal bovine serum (FBS; GeminiBio), 1X nucleosides (Sigma-Aldrich), 1000 U/mL leukemia inhibitory factor (LIF; Millipore), and 100 µM beta-mercaptoethanol (BME; Gibco). Cells were passaged every 2 days and seeded at a density of 1.75 x 10^6^ cells per flask in fresh complete medium.

### Preparing Embryoid Bodies (EBs)

Polydimethylsiloxane (PDMS) microwell inserts (400 x 400 µm inverted pyramids) were used to form 3D embryoid body (EB) aggregates of mESCs. Inserts were sterilized by sonicating twice in 100% isopropyl alcohol, placed in fresh 24-well plates, washed again with 100% isopropyl alcohol, then rinsed three times in excess 1X Dulbecco’s phosphate-buffered saline (dPBS) followed by UV sterilization for 30 minutes.

Prior to seeding cells, 500 µL of DFK5 medium was added to each insert and centrifuged at 2000 x g for 5 minutes to remove bubbles. DFK5 consisted of a DMEM/F12 base supplemented with 5% Knockout Serum Replacement (KSR; Gibco), 50 µM non-essential amino acids (NEAA; Gibco), 100 µM BME, 1:100 100X Insulin-Transferrin-Selenium (ITS; Gibco), 1:200 1X Nucleosides. 160,000 cells were seeded into each microwell insert then centrifuged at 200 x g for 5 minutes to achieve approximately 130 cells per EB. EBs were allowed to form undisturbed for 2 days.

### Embedding Embryoid Bodies (EBs) into Hydrogels

Solutions of HA-Nor-ADH, HA-IKVAV, and HA-RGD were dissolved at 3% (w/v) in 1X dPBS, while HA-Ox and Alg-Ox were dissolved at 6% (w/v) in dPBS. In a 48-well plate, 250 µL of gelatin were added to control wells and 100 µL of HA-Nor-ADH, HA-IKVAV, or HA-RGD were added to their respective wells.

For each plate, 2 mL of EB suspension was collected from microwells, centrifuged at 200 x g for 5 minutes, and resuspended in 400 µL of DFK5 medium. Next, 50 µL of EBs were gently mixed with HA hydrogels. Finally, 50 µL of the HA-Ox or Alg-Ox were gently mixed into their respective wells, resulting in a final hydrogel volume of 200 µL at a 3% (w/v) concentration. To avoid EB breakage, P200 pipette tips were trimmed using a sterile blade to increase the bore diameter. The hydrogels were allowed to crosslink for 30 minutes before adding 400 µL of DFK5 medium supplemented with 10 nM retinoic acid (RA; Sigma-Aldrich) and 1 µM purmorphamine (Pur; Millipore). The embedded EBs were then cultured for 2 days.

### Differentiation and Maturation of V2a Interneurons

Cells were enriched for V2a interneurons following previously established protocols in mESCs.^37,40,41^ 2 days of culture, the medium was replaced with DFK5 medium supplemented with 10 nM RA, 1 µM Pur, and 5 µM *N*-[N-(3,5-difluorophenacetyl)-l-alanyl]-S-phenylglycine t-butyl ester (DAPT). DAPT was included to inhibit the Notch signaling pathway and initiate differentiation to V2a interneruons. The EBs were cultured for an additional 2 days.

On Day 6 of the protocol, the cells cultured on gelatin were trypsinized for 5 minutes at 37°C using 0.25% trypsin-EDTA and transferred to laminin-coated wells. To prepare these wells, 150 µL of 0.01% (v/v) poly-L-ornithine solution was added to each well and incubated at 37°C for 1 hour. The poly-L-ornithine was aspirated, and the wells were washed five times with 250 µL Tris-buffered-saline (TBS). 100 µL of 0.01 mg/mL laminin diluted in HEPES buffered saline was added to each well and incubated overnight at 4°C. Prior to plating cells, the laminin solution was aspirated and washed once with HEPES buffered saline.

Approximately 175,000 V2a interneurons were plated in maturation medium (DFK5NB) in each laminin-coated well. DFK5NB consisted of a 1:1 ratio of DFK5 and neurobasal media (Gibco), supplemented with 1X B-27 (Gibco), and 1X GultaMAX (Gibco). The differentiation medium in the hydrogel-containing wells was similarly exchanged for 400 µL of DFK5NB maturation medium on Day 6. All cultures were then maintained until Day 12 of the protocol, with replacement of 50% of the medium performed every 2 days between Day 6 and Day 12 to yield mature V2 interneurons.

### Live/Dead Assay

Live/dead assays were performed per manufacturer’s instructions. Briefly, 1 µM of Calcein AM and 4 µM of Ethidium Homodimer-1 (EthD-1) were mixed in 1X dPBS, 300 µL of solution were added to each well and incubated at 37°C for 1 hour protected from light. Cells were then imaged using the Zeiss AxioObserver.

### RNA Extraction and Purification

Cells were collected for RNA extraction by adding 500 µL of TRIzol reagent (ThermoFisher) to each well and incubating at room temperature for 5 minutes. Following incubation, 200 µL of chloroform were added, and tubes were vortexed for 15 seconds. The tubes were incubated on ice for 10 minutes followed by centrifugation at 12,000 x g for 15 minutes at 4°C to induce phase separation. 150 µL of the upper aqueous phase were collected, mixed with 250 µL of 70% ethanol, and purified using the RNeasy Mini Kit (Qiagen) according to the manufacturer’s instructions. To elute the RNA, 30 µL of RNase-free water was added directly to the column membrane and centrifuged for 1 minute at 23,000 x g. To maximize RNA yield, the eluate was pipetted back onto the spin column and centrifuged for additional minute.

### cDNA Synthesis

The purified RNA samples underwent cDNA conversion using the High-Capacity RNA-to-cDNA kit (Applied Biosystems). Briefly, 10 µL of RT Buffer and 1 µL of RT Enzyme Mix were added to an 8-tube PCR strip. For each sample, 80 ng of RNA was added, and the final reaction volume was brought up to 20 µL using Ultrapure DNase/RNase free water (Invitrogen). The reverse transcription reaction was performed in a thermocycler with an incubation at 37°C for 60 minutes, followed by enzyme inactivation at 95°C for 5 minutes.

### Quantitative Real-Time PCR (qPCR)

For qPCR, a master mix was prepared for each target gene (*Chx10, Slc17a6, Mapt, Ppia*) by adding 5 µL of PowerTrack SYBR Green Master Mix (Applied Biosystems), 1 µL of a 100 µM primer mix **(Table 2)**, and 3 µL of ultrapure RNase/DNase free water per reaction. Next, 9 µL of the respective master mix and 1 µL of cDNA were added to the wells of an optical 96-well reaction plate (Applied Biosystems). The plates were sealed, vortexed, and centrifuged to remove bubbles. qPCR was performed using the BioRad CFX96 Real-Time PCR Detection System with the following thermal cycling conditions: an initial denaturation step at 95°C for 2 minutes, followed by 40 cycles of denaturation at 95°C for 15 seconds and annealing/extension at 60°C for 1 minute. Following amplification, a continuous melt curve analysis was performed (95°C for 15 seconds, 60°C for 1 minute, followed by a temperature ramp to 95°C) to verify product specificity.

**Table 2.**
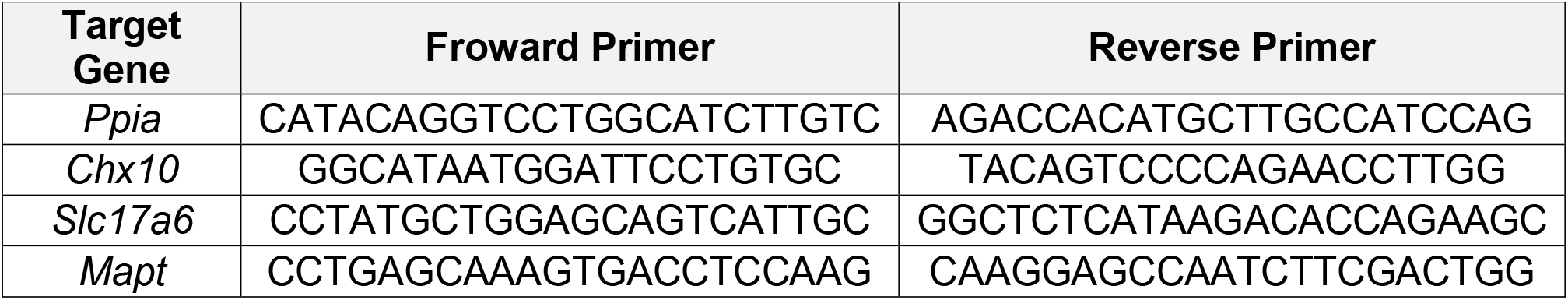
Table of primer pairs for qPCR listed 5’ to 3’.

**Table 3.** Table of antibodies used for immunostaining.

| Antibody | Species | Dilution | Catalog No. |
| --- | --- | --- | --- |
| Anti-βTubIII | Rabbit | 1:1000 | Biolegend 802001 |
| Anti-Rabbit 555 | Goat | 1:500 | Invitrogen A-21428 |

Relative gene expression was calculated by using the comparative cycle threshold (2^-ΔΔCq^) method. Target gene expression was normalized to *Ppia* as the housekeeping gene, and final gene expression levels were reported as log_2_ fold change relative to the control group (laminin).

### Immunocytochemistry

After 12 days of culture, cells were washed with 250 µL of 1X dPBS and fixed with 200 µL 4% (w/v) paraformaldehyde for 30 minutes at room temperature. The wells were then washed three times with 250 µL of 1X dPBS. To permeabilize cells, 200 µL of 0.1% (v/v) Triton X-100 in 1X PBS was added to each well and incubated at room temperature for 15 minutes. The wells were washed three times with 200 µL of washing buffer (0.1% (v/v) Tween-20 in 1X PBS).

To prevent non-specific binding, 200 µL of blocking buffer, consisting of 1% (w/v) bovine serum albumin (BSA), 22.5 mg/mL glycine, 5% (v/v) normal goat serum (NGS) and 0.1% (v/v) Tween-20 in PBS, were added to each well. The plates were incubated overnight at 4°C with shaking at 150 rpm.

The following day, wells were washed three times for 15 minutes each with 200 µL of washing buffer. Cells were then incubated with 200 µL of primary antibody solution, consisting of rabbit anti-βIII-tubulin (1:1000; BioLegend) in 1% (w/v) BSA and 2% (v/v) NGS in 1X PBS overnight at 4°C with shaking at 150 rpm.

After primary antibody incubation, the wells were washed three times for 15 minutes each with 200 µL of washing buffer. Next, 200 µL of secondary antibody solution containing goat anti-rabbit 555 (1:500; Invitrogen) and DAPI (1:1000; Invitrogen) in 1% (w/v) BSA and 2% (v/v) NGS in 1X PBS were added to each well. Plates were incubated overnight at 4°C with shaking at 150 rpm.

Following the secondary antibody incubation, the wells were washed three times with washing buffer with gentle agitation at 20 rpm for 30 minutes at room temperature. Finally, 200 µL of 1X PBS were added to each well, and the cells were imaged using the Zeiss Observer microscope.

### Live/Dead Image Analysis

All image analysis was performed with ImageJ. Images were converted to 8-bit image type, and z stacks were projected. Channels were split for Calcein AM (green) and EthD-1 (red). The background was subtracted using a rolling ball radius of 600 pixels, and image contrast was enhanced for saturated pixels of 0.35%. An Ostu threshold was performed and adjusted to include aggregates. Binary images were then cleaned by filling holes, and open commands were used to remove debris. The area and integrated density were measured, and particles larger than 100 µm^2^ were analyzed. The percentage viability was calculated using **equation 3**. Brightness and contrast were adjusted using the set display range and applied to all images. The display range for the green channel was 501-63888, and the display range for the red channel was 230-17722.

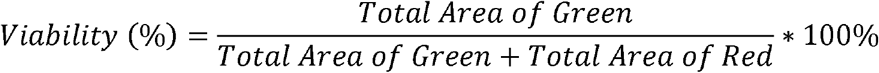

### Neurite Length Analysis

Image brightness and contrast were adjusted using the set display range and applied to all images. The display range for the blue channel was 16-40, and the display range for the red channel was 50-86. Neurite lengths from the aggregate were manually measured using the neuroanatomy plug in and the simple neurite tracer software in ImageJ.

## Results

### Polymer Modifications for the Synthesis of Hydrazone Crosslinked HA-Alg Hydrogels

To form hydrazone-crosslinked HA-Alg hydrogels (Fig. 1), alginate was oxidized resulting in a ring break, exposing aldehydes for crosslinking (Alg-Ox). A hydroxylamine hydrochloride titration was performed to calculate average percent degree of modification (%DOM) or the percentage of aldehydes per chain of alginate, which was 61.26 ± 8.82% with an average percent yield of 72.65 ± 10.93%. HA was then modified with norbornene functional groups as previously described and optimized for a low degree of modification to favor the downstream peptide conjugation.^50,52^ The average %DOM or the number of norbornene functional groups per chain of hyaluronic acid was 4.75 ± 2.21% with an average percent yield of 42.77 ± 1.93%. HA-Nor was then functionalized with adipic acid dihydrazide (HA-Nor-ADH) for crosslinking with Alg-Ox. An average %DOM of adipic acid dihydrazide groups per chain of HA determined by 1H NMR was 37.50 ± 5.71% with an average percent yield of 65.19 ± 5.29 %. HA-Nor ADH was then used to conjugate cysteine terminal peptides, IKVAV and RGD using a photoinitiated thiol-ene click chemistry reaction (HA-IKVAV or HA-RGD). The average concentration of peptides conjugated to the backbone of HA was determined to detect using an Ellman’s assay which detects the disappearance of free thiol groups before and after UV exposure and conjugation. The average concentration of IKVAV on the polymer was determined to be 12.69 ± 6.10 nmol/mg HA, and the average concentration of RGD on the polymer was determined to be 14.27 ± 7.55 nmol/mg HA. A summary of polymer modifications quantifications can be found in **Tabel S1**.

**Figure 1.**
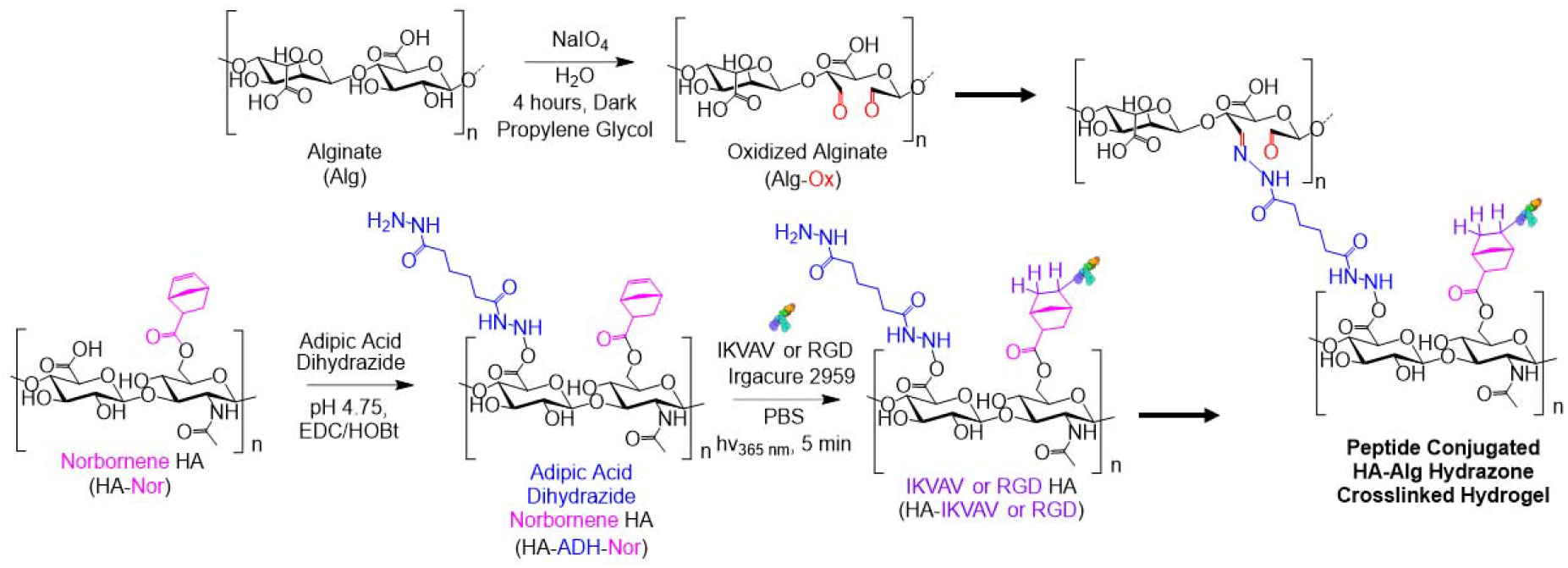
Polymer modifications for the fabrication of HA-Alg hydrogels. Alginate was oxidized to reveal aldehyde groups for crosslinking (Alg-OX). An EDC/HoBt coupling reaction was performed with HA-Nor to form HA functionalized with adipic acid dihydrazide and norbornene (HA-Nor-ADH). A photoinitiated reaction between HA-Nor-ADH and the terminal cysteine on the IKVAV or RGD peptide was then performed with UV light at a wavelength of 365 nm. The terminal products of these reactions, HA-Nor-ADH and Alg-Ox, were mixed at different polymer concentrations to produce hydrazone crosslinked hyaluronic acid-alginate (HA-Alg) hydrogels.

### HA-Alg Hydrogels Exhibit a Range of Physicochemical Properties

HA and alginate were mixed at different polymer concentrations (1-3% (w/v)) to form hydrogels. We confirmed hydrogel injectability by passing through a needle and the physiochemical properties of the HA-Alg hydrogels were evaluated to determine the optimal hydrogel formulation for neuronal culture. Our target gelation time was between 1 to 5 minutes to produce a workable hydrogel for embedding mESCs for V2a interneuron differentiation and maturation. Increasing the polymer concentrations of the hydrogel decreased the gelation time, likely due to the increase of functional groups available for crosslinking. All hydrogels gelled within 5 minutes except for the HA_1.0_Alg_1.0_, HA_1.0_Alg_2.0_, and HA_2.0_Alg_1.0_ hydrogels with lower polymer concentrations (Fig. 2A). We targeted a hydrogel stiffness that would match the stiffness of CNS tissue (100-1000 Pa). We observed an overall increase in compressive modulus with increasing polymer concentrations (Fig. 2B). The HA_3.0_Alg_3.0_ hydrogel was the only hydrogel to display a compressive modulus above 100 Pa. Lastly, we measured the mass change of the hydrogels in both PBS and aCSF over 28 days to observe swelling and degradation over time (Fig. 2C-D). We targeted a stable hydrogel, defined as a mass change of no more than ± 25% of its original mass, over 28 days to enable long-term neuronal culture. Hydrogel stability in both PBS and aCSF decreased as polymer concentrations deviated from a 1:1 stoichiometric ratio. Additionally, hydrogel formulations with higher alginate concentrations were less stable in aCSF compared to PBS due the increased concentration of ions in the solution.

**Figure 2.**
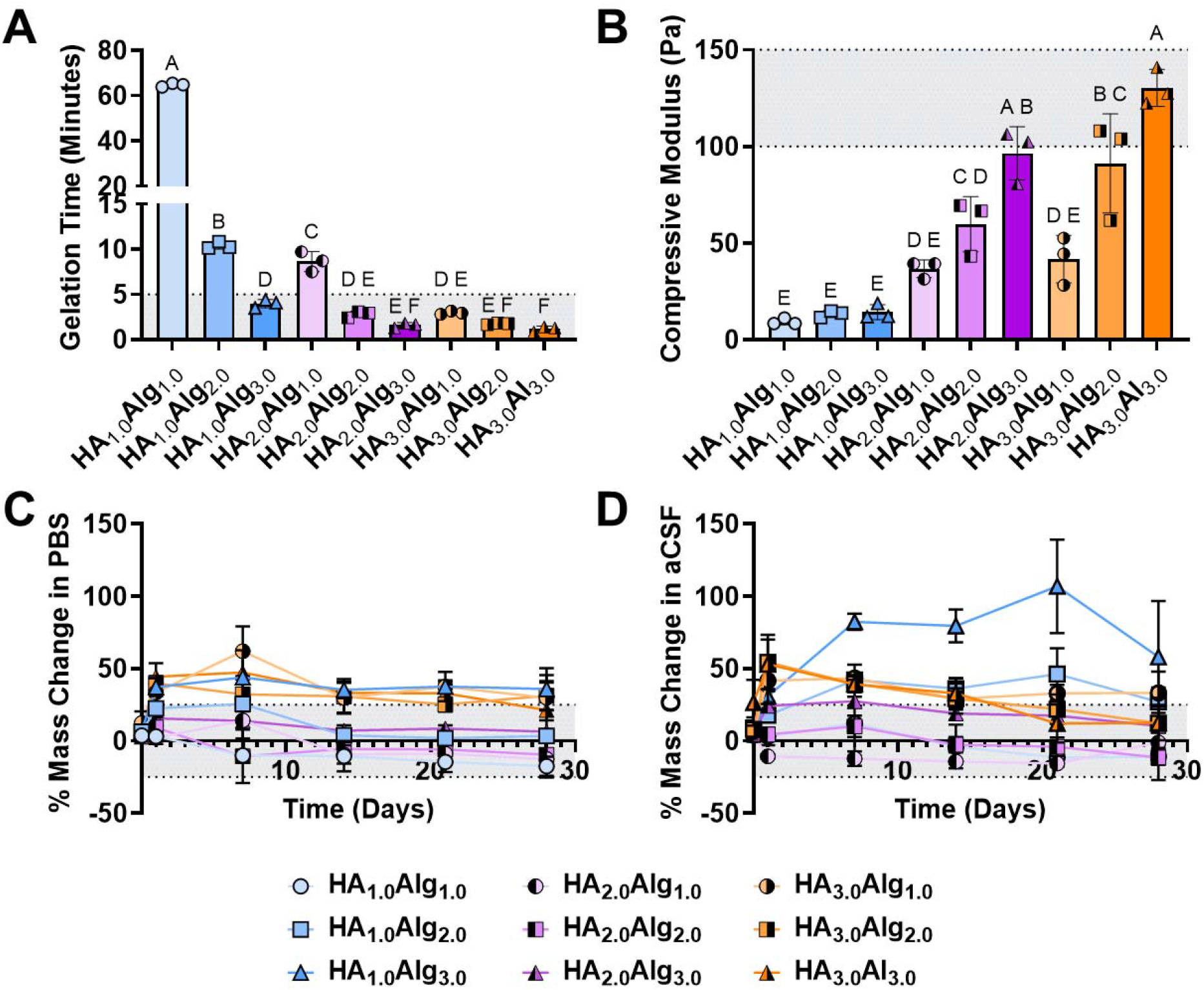
Physiochemical properties of HA-Alg hydrogels. HA-Alg hydrogels with various polymer concentrations of HA and alginate denoted by subscripts (HA_% w/v_Alg_% w/v_). Shaded regions indicate desired hydrogel properties. A) The gelation time of HA-Alg hydrogels decreased with increasing polymer concentration. B) The compressive modulus of HA-Alg hydrogels increased with increasing polymer concentration. C) Mass change of HA-Alg hydrogels in phosphate-buffered saline (PBS) over 28 days. D) Mass change of HA-Alg hydrogels in artificial cerebrospinal fluid (aCSF) over 28 days. One-way ANOVA, post hoc Tukey’s multiple comparisons was performed for individual replicates. Statistics are summarized using the Compact Letter Display, where differing letters (A, B) represent groups that are statistically different (p < 0.05) and the same letter represents groups with no statistical differences (p > 0.05) for three technical replicates (n=3).

We determined that the hydrogel formulation HA_3.0_Alg_3.0_ displayed physicochemical properties that best aligned with our desired properties for neural cell culture. This hydrogel exhibited an average gelation time of 1. 2 ± 0.3 minutes and compressive modulus of 130.3 ± 9.6 Pa. Although the HA_3.0_Alg_3.0_ lost more than 25% of its original volume in the first 14 days in both PBS and aCSF, the hydrogel stabilized within 25% of its original mass by day 28. Conjugating IKVAV and RGD peptides to HA-Nor-ADH to form peptide-containing hydrogels resulted in approximately 0.38 ± 0.18 mM of IKVAV per hydrogel and 0.43 ± 0.23 mM of RGD per hydrogel which falls within the concentration needed for neural cell adhesion.^53,54^

### HA-Alg Hydrogels Support V2a Interneuron Viability

To evaluate the ability of the HA-Alg hydrogels to support neuronal culture, V2a interneurons were differentiated from mESCs and cultured over 12 days in HA-Alg hydrogels and compared to cells cultured in previously established HA-HA hydrogels and 2D gelatin/laminin-coated control wells.^37,40,41^ Following 2 days of forced aggregation, EBs were mixed with either unmodified or peptide-modified HA-ADH, either HA-Ox or Alg-Ox was added, and the hydrogels were crosslinked for 30 minutes. After 12 days of culture, quantitative live/dead analysis demonstrated that the incorporation of the laminin-derived IKVAV peptide was essential for sustaining 3D cell viability at levels comparable to standard 2D laminin controls (Fig 3 A-B). While viability did not differ between any of the hydrogels themselves, cells embedded in the unmodified HA+Alg hydrogels (54.9 ± 2.6%) and RGD-modified HA-RGD+HA and HA-RGD+Alg hydrogels (50.0 ± 7.7% and 52.7 ± 1.4%, respectively) exhibited significantly lower viability than cells in the 2D laminin control (91.4 ± 6.0%). In contrast, IKVAV-modified HA-IKVAV+HA and HA-IKVAV+Alg hydrogels supported cell survival, yielding viabilities (77.1 ± 7.2% and 72.4 ± 20.9%) that were comparable to the 2D laminin control.

**Figure 3.**
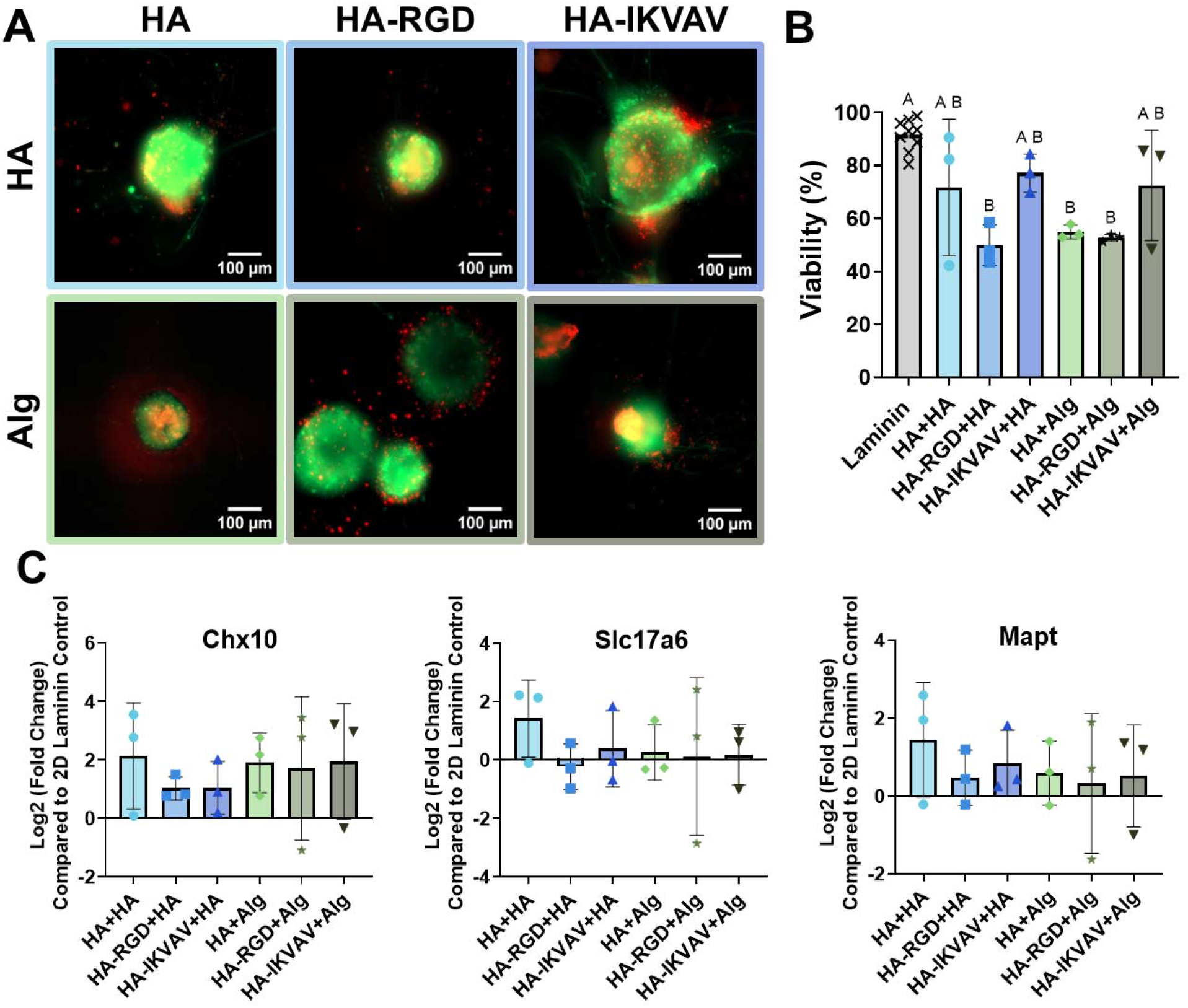
V2a interneurons remain viable and express neural marker expression in HA-Alg and HA-HA hydrogels. A.) Representative live/dead fluorescent images of cells encapsulated within hydrogels after 12 days in culture. Live cells are stained green (Calcein-AM), and dead cells are stained red (EthD-1). Scale bars = 100 µm. B.) Quantification of cell viability expressed as the percentage of live cells over total cells. Data are presented as mean ± standard deviation. One-way ANOVA, post hoc Tukey’s multiple comparisons was performed for individual biological replicates (N = 3). Statistics are summarized using the Compact Letter Display, where differing letters (A, B) represent groups which are statistically different (p < 0.05) and the same letter represents groups with no statistical differences (p > 0.05). C.) Relative gene expression (Log2 Fold Change) of neural differentiation markers *Chx10, Mapt*, and *Slc17a6* quantified via RT-qPCR. Bars represent the mean Log2 fold change relative to the 2D laminin control, and error bars represent the standard error of the mean (SEM). Individual datapoints represent independent biological replicates (N = 3). No statistically significant differences were found between hydrogel formulations (p > 0.05) using a one-way ANOVA with Tukey’s post-hoc test.

### Evaluation of V2a Interneuron Differentiation in HA-Alg Hydrogels

To assess if the culture had been successfully enriched for V2a interneurons after 12 days, quantitative real-time PCR (qPCR) was performed. Gene expression profiles of cells embedded in all hydrogel formulations were compared to those of cells cultured on standard 2D laminin coatings (Fig. 3C). Primer specificity and reaction efficiencies were confirmed via melt curve analysis and amplification plots (Fig. S3, S4), and the stable expression of the endogenous control gene *Ppia* was verified across all samples.

Overall, embedding cells within the hydrogels promoted upregulation of V2a interneuron markers (*Chx10*) and general neuronal maturation markers (*Mapt* and *Slc17a6*) compared to the 2D laminin control (Fig. 3C). Expression of *Chx10*, a transcription factor expressed in V2a interneurons, was upregulated across all hydrogel formulations. Interestingly, despite their restricted morphological development, unmodified hydrogel formulations demonstrated some of the highest relative expression levels of *Chx10*. To assess the maturation of the differentiated neurons, the expression of *Mapt*, a general marker of mature neurons, and *Slc17a6*, the gene that encodes for VGlut2, a marker of excitatory glutamatergic neurons, were evaluated. Similar to the *Chx10* profiles, *Mapt* and *Slc17a6* exhibited positive fold changes across most of the hydrogel conditions compared to laminin.

Importantly, all hydrogel formulations supported comparable expression of *Chx10 and Slc17a6* compared to 2D laminin controls, confirming successful enrichment of V2a interneurons. The comparable gene expression profiles observed across the different hydrogel formulations indicate that the small molecules driving differentiation were efficiently diffusing through the 3D polymer networks, regardless of the polymer type, crosslinking chemistry, and peptide modification. These transcriptional profiles further suggest that early lineage specification was primarily driven by soluble media cues rather than matrix-bound adhesive signals.

### IKVAV-modified HA-Alg Hydrogels Enhance V2a Interneuron Neurite Outgrowth

The surrounding hydrogel network heavily influenced the morphology and structural maturation of the cells in 3D. To characterize this development, we evaluated EB morphology alongside immunostaining for the neuron-specific microtubule marker βIII-Tubulin (TUBB3) after 12 days of culture. As expected, cells cultured on 2D laminin controls exhibited characteristic planar cellular adhesion and dispersed TUBB3 positive neurite networks (Fig. S5). In 3D cultures, the biochemical makeup of the microenvironment dictated the structural fate of the V2a interneurons (Fig. 4A, S5). EBs encapsulated within the unmodified hydrogels (HA+HA and HA+Alg) formed dense, DAPI-positive spheroids that maintained tightly restricted morphologies. These structures were characterized by distinct smooth borders and an absence of outward neurite extension. Interestingly, modifying hydrogels with the fibronectin-derived RGD peptide (HA-RGD+HA and HA-RGD+Alg) failed to overcome this physical confinement. TUBB3 staining indicated a lack of macroscopic cytoskeletal extensions with minimal expression restricted to the EB surface. While a small fraction of EBs within these unmodified and RGD functionalized environments occasionally exhibited small projections, overall structural development was negligible.

**Figure 4.**
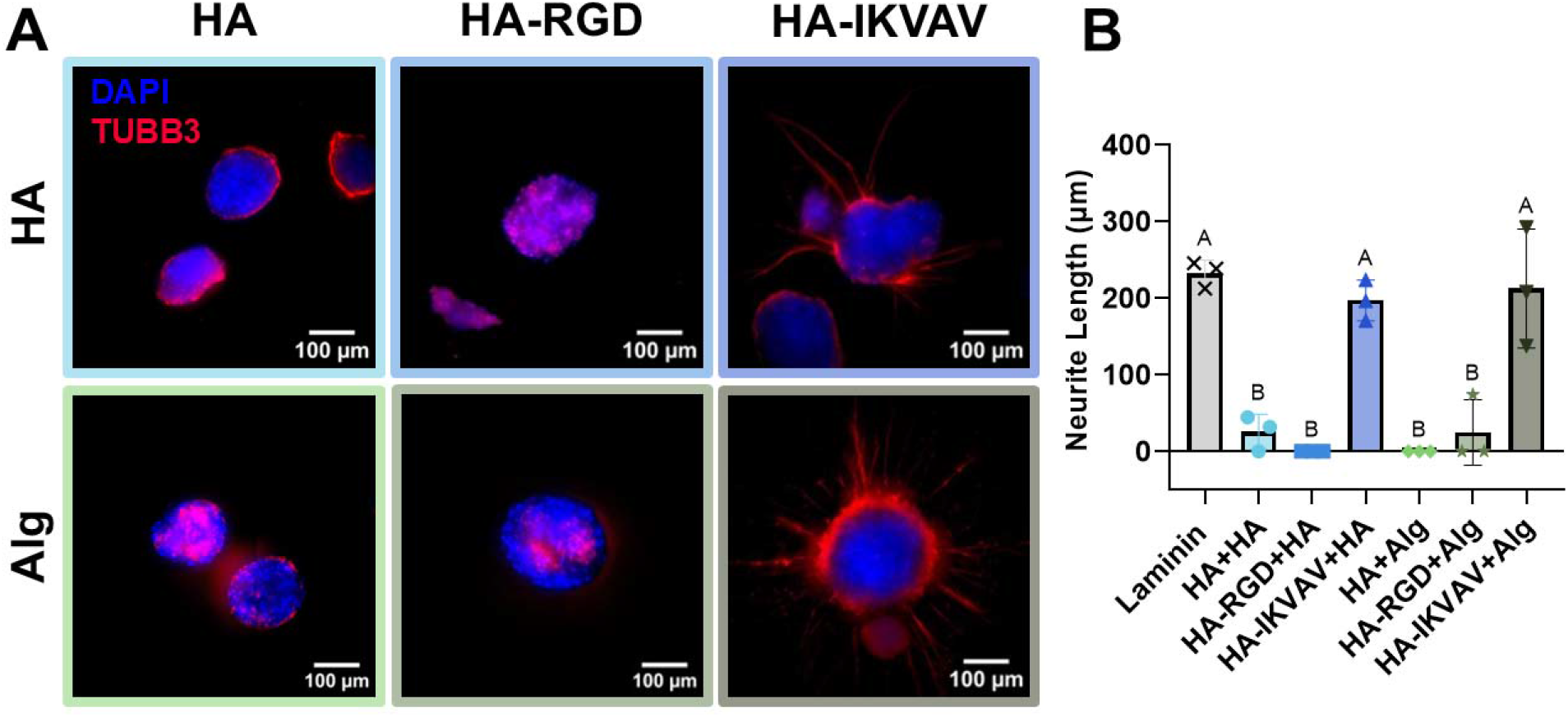
Immunostaining assessing V2a interneuron maturation and neurite length. A.) Representative maximum intensity projections of V2a interneurons cultured for 12 days in HA-HA and HA-Alg crosslinked hydrogels (unmodified, RGD, and IKVAV). Cells were immunostained for the neuronal microtubule marker βIII-Tubulin (TUBB3, red) and nuclei (DAPI, blue). Scale bars = 100 µm. B.) Quantification of average neurite length (µm) extending from the embryoid bodies in various hydrogel formulations and the 2D laminin control. One-way ANOVA, post hoc Tukey’s multiple comparisons was performed for individual biological replicates with three technical replicates (N = 3, n=3). Error bars represent the standard error of the mean (SEM). Statistics are summarized using the Compact Letter Display, where differing letters (A, B) represent groups which are statistically different (p < 0.05) and the same letter represents groups with no statistical differences (p > 0.05).

Conversely, functionalization with the laminin-derived-peptide IKVAV (HA-IKVAV+HA and HA-IKVAV+Alg) established a permissive microenvironment that facilitated extensive structural development. EBs within IKVAV-modified hydrogels demonstrated clear morphological maturation, marked by TUBB3 positive neurites projecting radially into the surrounding 3D hydrogel space. These robust microtubule networks demonstrate that IKVAV not only permits physical cell spreading but actively supports the cytoskeletal assembly required for mature neuronal process extension. While the presence and length of extending neurites naturally exhibited some heterogeneity within samples, outgrowth within the IKVAV-containing hydrogels was overwhelmingly consistent. Quantitative analysis of neurite length corroborated these visual observations (Fig. 4B). IKVAV-modified hydrogels supported substantial neurite outgrowth, averaging approximately 197.2 ± 53.5 µm for HA-IKVAV+HA and 212.4 ± 75.2 µm for HA-IKVAV+Alg hydrogels that was comparable standard 2D laminin controls (232.3 ± 25.6 µm) where robust neurite extension would be expected. Conversely, the unmodified and RGD-modified hydrogels yielded negligible extensions. This morphological divergence demonstrates that incorporating targeted laminin-derived peptides like IKVAV effectively breaks the physical restrictions around the EBs and drives the maturation and neurite outgrowth of V2a interneurons in 3D. Taken together with the gene expression profiles, while chemical differentiation proceeded efficiently across all groups, the physical architecture and biochemical cues provided for the IKVAV-modified hydrogels was critical for structural maturation.

## Discussion

The successful maturation of neurons within defined 3D hydrogel microenvironments is a critical step toward developing effective cell-based regenerative therapies for SCI. In this study, we developed hydrazone crosslinked HA-Alg hydrogels modified with fibronectin- and laminin-derived peptides (RGD and IKVAV, respectively) and evaluated their capacity to support the viability, lineage specification, and morphological development of mESC-derived V2a interneurons.

A primary challenge in neural tissue engineering is designing scaffolds that mimic the environment of the CNS without sacrificing 3D architecture. By optimizing the polymer concentration to the HA_3.0_Alg_3.0_ formulation we achieved a compressive modulus of 130.3 ± 9.6, aligning directly with the properties of native CNS tissue (100-1000 Pa). Previous literature has demonstrated that excessive stiffness drives neural progenitor cells toward glial lineage and exacerbate astrogliosis, whereas compliant matrices are permissive to neurogenesis and robust neurite outgrowth.^55^ In addition to matching target stiffness, the gelation kinetics of our HA-Alg system were optimized to address the practical challenges of embedding cells in 3D. Slower-gelling systems frequently suffer from gravity-induced cell sedimentation before crosslinking is complete, leading to heterogeneous cell distributions, localized nutrient deprivation, and inconsistent cell-matrix interactions. We observed that the HA_3.0_Alg_3.0_ formulation gels within 1. 2 ± 0.3 minutes, rapid enough to ensure uniform 3D suspension.^56^

The most significant physicochemical advantage of our system is its long-term stability in physiological fluids. Conventional alginate hydrogels rely on transient ionic crosslinks, which rapidly dissociate in sodium-rich or chelator-containing media due to ion exchange.^57^ This premature dissolution limits their utility in neural cultures, as neurons require extended timelines for functional maturation. By using covalent hydrazone bonds between oxidized alginate and ADH-modified HA, our hydrogels overcame this limitation, stabilizing within 25% of their original mass over 28 days in ion-rich aCSF.

We compared HA-Alg hydrogels to a previously established HA-HA hydrogel to specifically evaluate the effects of alginate on cell viability and maturation. While HA is a ubiquitous component of the neural ECM, our results demonstrated that an HA-Alg hybrid system is equally cytocompatible compared to HA-HA hydrogels. This finding is significant as it validates the use of alginate as a tunable and clinically relevant copolymer for neural cell culture that does not compromise the ability of the scaffold to support neural differentiation and neurite outgrowth.

Our findings reveal a notable divergence between differentiation and morphological maturation within these 3D environments. While V2a interneuron differentiation was efficiently driven by soluble cues across hydrogel conditions, we observed that structural maturation and neurite outgrowth required specific ECM-derived peptides (Fig. 4A). Specifically, our findings highlight that, while both the HA-Alg and HA-HA backbones provided a stable structural system for supporting cell viability, the presence of laminin-derived IKVAV was the essential driver for neurite extension, thereby dictating functional cellular behavior.

Gene expression profiling revealed that embedding within both peptide-modified and unmodified hydrogels supported the upregulation of the V2a interneuron transcription factor *Chx10* and mature neuronal markers (*Mapt, Slc17a6*) (Fig. 3C). The success of this differentiation across all hydrogel formulations indicates that the small molecules added for differentiation were equally effective across the different hydrogel networks and were not hindered by the crosslinking chemistry or presence of peptides. The successful upregulation of *Chx10* is a critical benchmark for V2a interneuron differentiation, as *Chx10*^*+*^ V2a interneurons play an integral role in driving the excitatory pathways of motor neurons and are essential for locomotor initiation. In fact, recent studies have demonstrated that V2a interneurons are explicitly required for the recovery of critical motor functions following cervical SCI, making them a primary therapeutic target for regenerative medicine.^34,39,58,59^

Interestingly, the morphology of V2a interneurons varied drastically across hydrogel formulations. While unmodified hydrogels and hydrogels modified with RGD exhibited tightly confined EB morphology and lacked meaningful neurite projections, IKVAV-modified hydrogels facilitated radial neurite extension (Fig. 4A, S5). Furthermore, immunostaining for TUBB3 revealed dense, radially aligned microtubule networks (Fig. 4A, S5). This structural divergence highlights that, while the basic physical properties of the hydrogels may permit cell survival, tissue-specific biochemical cues were required to drive structural maturation. Universal cell-adhesive motifs like RGD have been shown to successfully support neural differentiation and neurite extension in specific defined 3D microenvironments, including HA and polyethylene glycol-based hydrogels.^60,61^ However, although few V2a interneurons exhibited spontaneous neurite extension in RGD-modified hydrogels similar to unmodified HA hydrogels, the morphological maturation within RGD networks proved highly variable and less robust than IKVAV-functionalized hydrogels. These findings reflect established *in vitro* standards, as laminin is often used in neural differentiation protocols. Furthermore, laminin plays a central role in the native CNS ECM, targeting specific integrin pathways that are essential for neural maturation. By integrating the IKVAV peptide into our 3D scaffold, we successfully recapitulated the biochemical cues traditionally provided by 2D laminin coatings, thereby better mimicking the native CNS environment.

Notably, our peptide conjugation strategy yielded relatively low peptide concentrations compared to conventional hydrogels, resulting in approximately 0.38 ± 0.18 mM IKVAV and 0.43 ± 0.23 mM RGD per hydrogel compared to 2-6 mM observed in literature.^62–64^ This quantification offers a compelling explanation for the divergent morphological outcomes, which are likely driven by a combination of the low viability observed in RGD-modified hydrogels and the context-dependent biphasic influence of generic integrin-binding peptides. For example, one study demonstrated that low densities of RGD may provide baseline traction required for growth, but higher ligand densities can rapidly establish excessively rigid focal adhesions.^65^ These strong adhesions can anchor the growth cone to the matrix and restrict motility, preventing neurite outgrowth. Consequently, minor batch-to-batch variations in macroscopic hydrogel properties or localized peptide presentation can shift an RGD matrix from a permissive state to a restrictive one. However, given that our peptide conjugation yielded relatively low concentrations, it is more likely that our RGD hydrogels lacked the ligand density required to reach the critical threshold for neural adhesion. This aligns with findings from neurospheres seeded on top of similar HA-RGD-modified hydrogels, which showed that neurite density is dependent on precise RGD concentration.^61^ In contrast, IKVAV mimics the native laminin-rich ECM of the central nervous system, engaging neural specific receptors to promote true microtubule assembly, possibly requiring a lower concentration of conjugated peptide for a robust morphological response. While fibronectin-binding integrins are expressed in early stages of stem/progenitor cell neuronal differentiation, they are reported to be downregulated as neurons mature,^66^ and late-stage neurons predominantly express laminin-binding integrins (α1β1 and α6β1).^67^ These differences in integrin expression further support the hypothesis that higher concentrations of RGD are needed to activate cell adhesion while lower concentrations of IKVAV may be needed to elicit similar responses.

Native brain and spinal cord ECM heavily rely on complex proteoglycans and structural proteins to regulate the adhesion of neurons and drive subsequent neurite extension.^68^ Our results align with recent studies demonstrating that polymers such as alginate or HA lack the requisite motifs to engage neural integrins, thereby acting as physical barriers rather than permissive scaffolds unless modified. For instance, alginate hydrogels functionalized with laminin and gelatin significantly enhanced the neurite outgrowth and migration of human induced pluripotent stem cell derived neurospheres, suggesting that ligand-receptor interactions are required to trigger the cytoskeletal remodeling necessary for axonal extension.^69^ Similarly, hydrogels containing alginate and collagen supported robust neural network formation.^70^ The success of these hybrid systems is largely attributed to the fact that gelatin and collagen naturally present a high density of cell-adhesive sequences, including RGD, which provide the necessary biochemical anchors for neural attachment. However, when these general motifs support broad neural network formation, V2a interneurons appear to require more specialized signaling for functional development. Specifically, one study found that the incorporation of astrocyte-derived protoplasmic ECM into HA hydrogels promoted V2a interneuron process extension, with the ECM abundant in fibronectin, collagen, perlecan, and laminin.^44,71^ In another study, laminin was required to support the viability of V2a interneurons embedded in HA hydrogels and treated with neurotrophin-3.^8^ These findings supporting our observation that V2a interneurons are highly sensitive to their microenvironment and require laminin-derived ECM recognition cues for their maturation.

Our use of precise peptide conjugation offers a defined approach to engaging the cell’s mechanotransducive machinery. By utilizing IKVAV peptides, we successfully bypassed the restrictive physical boundaries of the hydrogel, allowing mESC-derived interneurons to achieve maturation. The heterogeneity in neurite length observed within these functionalized groups likely reflects the variability of mESC differentiation kinetics and intra-EB spatial positioning, which could be further optimized in future studies by tuning the peptide density or the degradation rate of the polymer network.

## Conclusion

We successfully engineered an injectable, hydrazone-crosslinked HA-Alg hydrogel designed to meet the complex physicochemical and biochemical requirements of neuronal cell culture with eventual applications in CNS tissue repair. We discovered a fundamental decoupling of biochemical differentiation and morphological maturation of V2a interneurons within 3D microenvironments. While small molecule for neuronal differentiation efficiently permeated all hydrogel formulations to drive the robust transcriptional differentiation of *Chx10*^*+*^ V2a interneurons, structural maturation required biochemical cell-adhesive cues. Modification of the hydrogels with the laminin-derived IKVAV peptide transformed the matrix into a highly permissive scaffold that maintained cell viability and drove extensive neurite growth. Furthermore, the HA-Alg composite hydrogel outperformed the similarly crosslinked HA-HA network in facilitating this structural maturation, highlighting the advantages of using blended polymer systems. Ultimately, the HA-IKVAV+Alg hydrogel represents a highly tunable, biomimetic platform capable of overcoming the physical restrictions of encapsulating stem cell aggregates in hydrogels. By successfully supporting the survival, differentiation, and structural maturation of therapeutically critical V2a interneurons, this injectable biomaterial offers a promising, minimally invasive strategy to transplant cells to promote functional regeneration following SCI in the future.

## Acknowledgements

We are grateful for funding from the National Institutes of Health (R21-EB032112), Wu Tsai Human Performance Alliance, Joe and Clara Tsai Foundation, and Donald E. and Delia B. Baxter Foundation as well as support from the Lary Simpson Professorship to M.H.H. A.N.G. is supported by the Giustina Knight Campus Fellowship Program, A.K.C. was supported by a Wu Tsai Performance Undergraduate Fellowship, and J.D.K. was supported by the Knight Campus Undergraduate Scholars program. We are also grateful to Chandler Asnes for technical support on image analysis.

## Data Availability Statement

The data supporting this article have been included as part of the Supplementary Information.

