## Supplemental Information for "Hyaluronic Acid-Alginate Hydrazone Crosslinked Hydrogels Support the Generation and Maturation of V2a Interneurons"

**Title**:

**Supplemental Information**

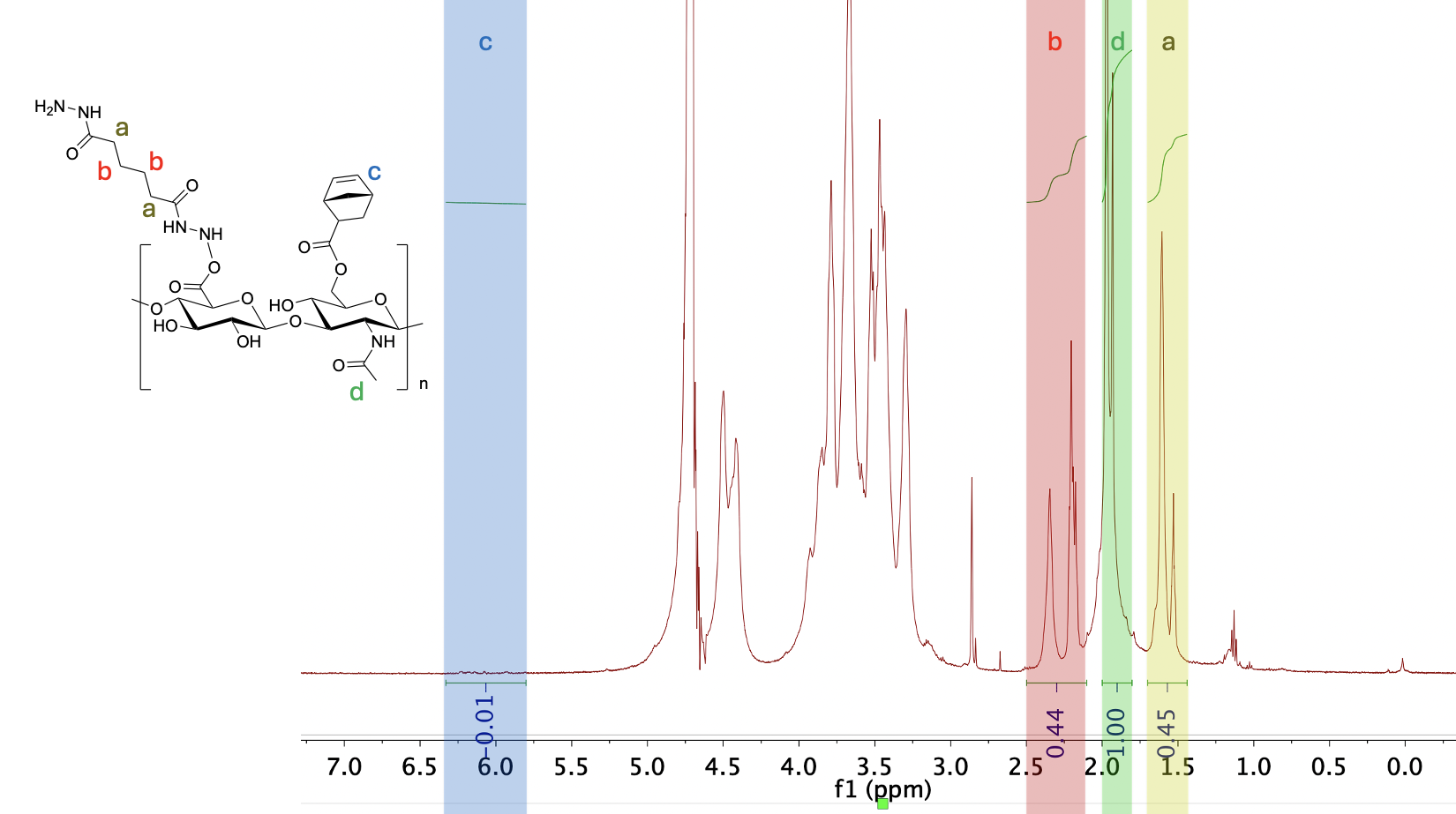

**Figure S1. ^1^H-NMR spectra of HA-Nor-ADH.** Norbornene (HA-Nor) and adipic acid dihyrazide (HA-Nor-ADH) modifications of hyaluronic acid were confirmed by ^1^H-NMR spectroscopy. Degree of modification of ADH was determined by peaks found at 1.44- 1.7 ppm (a) and 2.1-2.5 ppm (b) (8H). Degree of modification of norbornene was determined by peaks found at 5.7 -6.3ppm (2H). Both peaks were normalized to the n-acetyl group on HA found at 1.8-2 ppm (3H, d).

**Tabel S1**. Summary of polymer modification quantification.

|  | **Degree of Modification (%)** | **Percent Yield** |
| --- | --- | --- |
| **Alg-Ox** | 61.26 ± 8.82 | 72.65 ± 10.93 |
| **HA-Nor** | 4.75 ± 2.21 | 42.77 ± 1.93 |
| **HA-Nor-ADH** | 37.50 ± 5.71 | 65.19 ± 5.29 |
|  | **Peptide Concentration (nmol/mg HA)** |  |
| **HA-RGD** | 14.27 ± 7.55 | - |
| **HA-IKVAV** | 12.69 ± 6.10 | - |

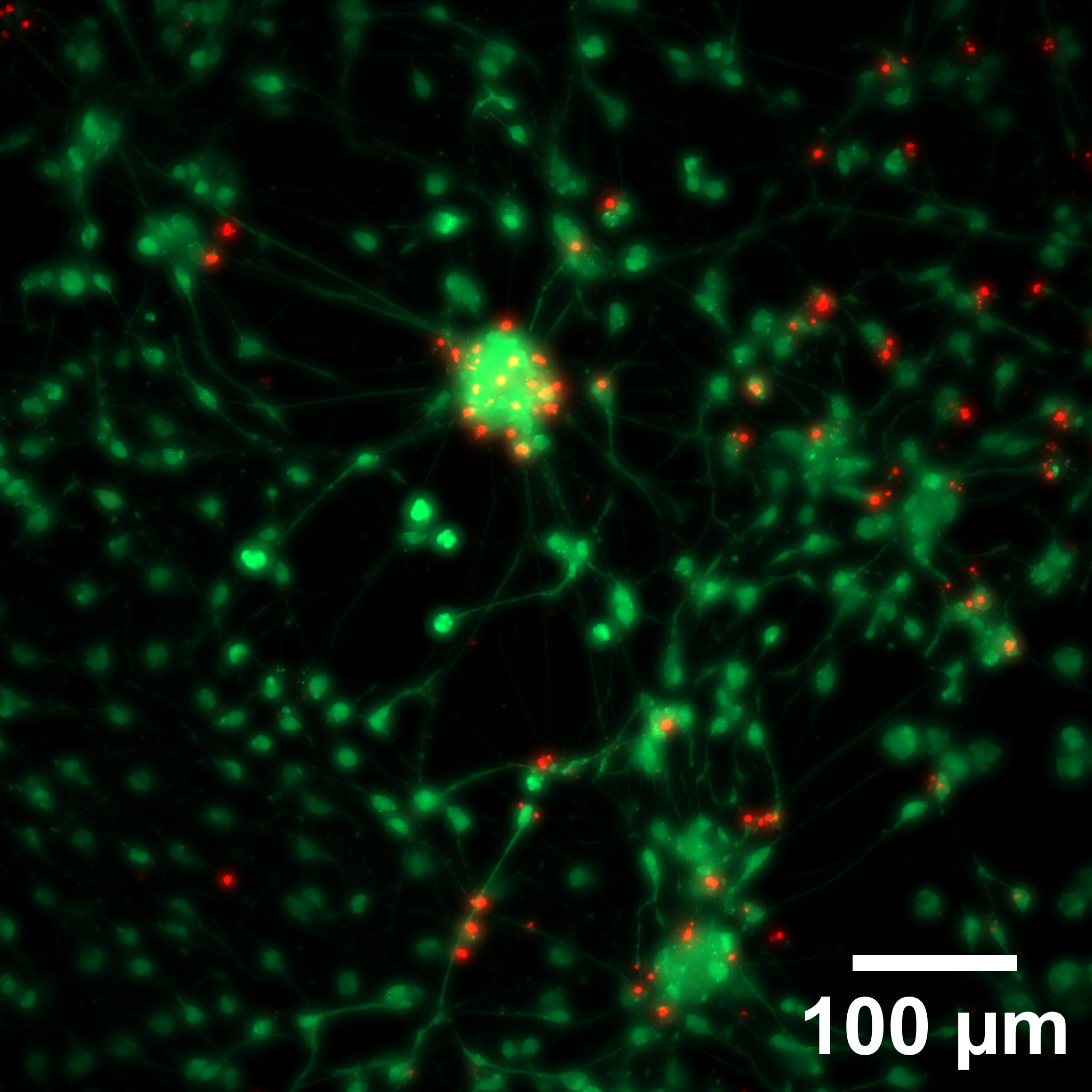

**Figure S2. Represent live/dead staining of cells cultured on 2D laminin coating.** Live cells are stained green (Calcein-AM), and dead cells are stained red (EthD-1). Scale bar = 100 µm.
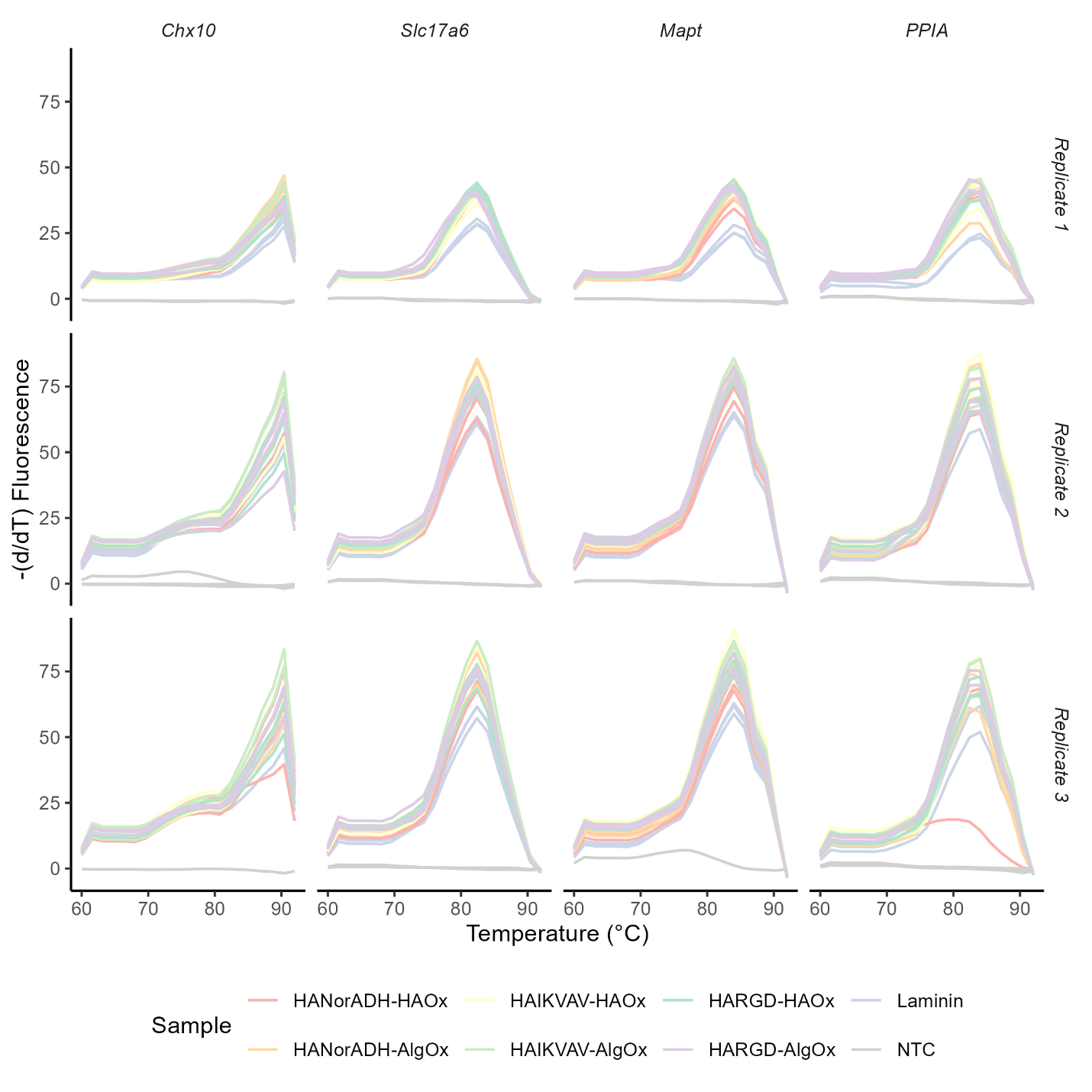
**Figure S3. Melt curve analysis for qPCR.** Melt curves represented as the negative derivative of fluorescence with respect to temperature for target genes. The presence of a single peak for each gene confirms primer specificity and absence of non-specific amplification. Columns correspond to a specific gene target, and rows represent independent biological replicates.

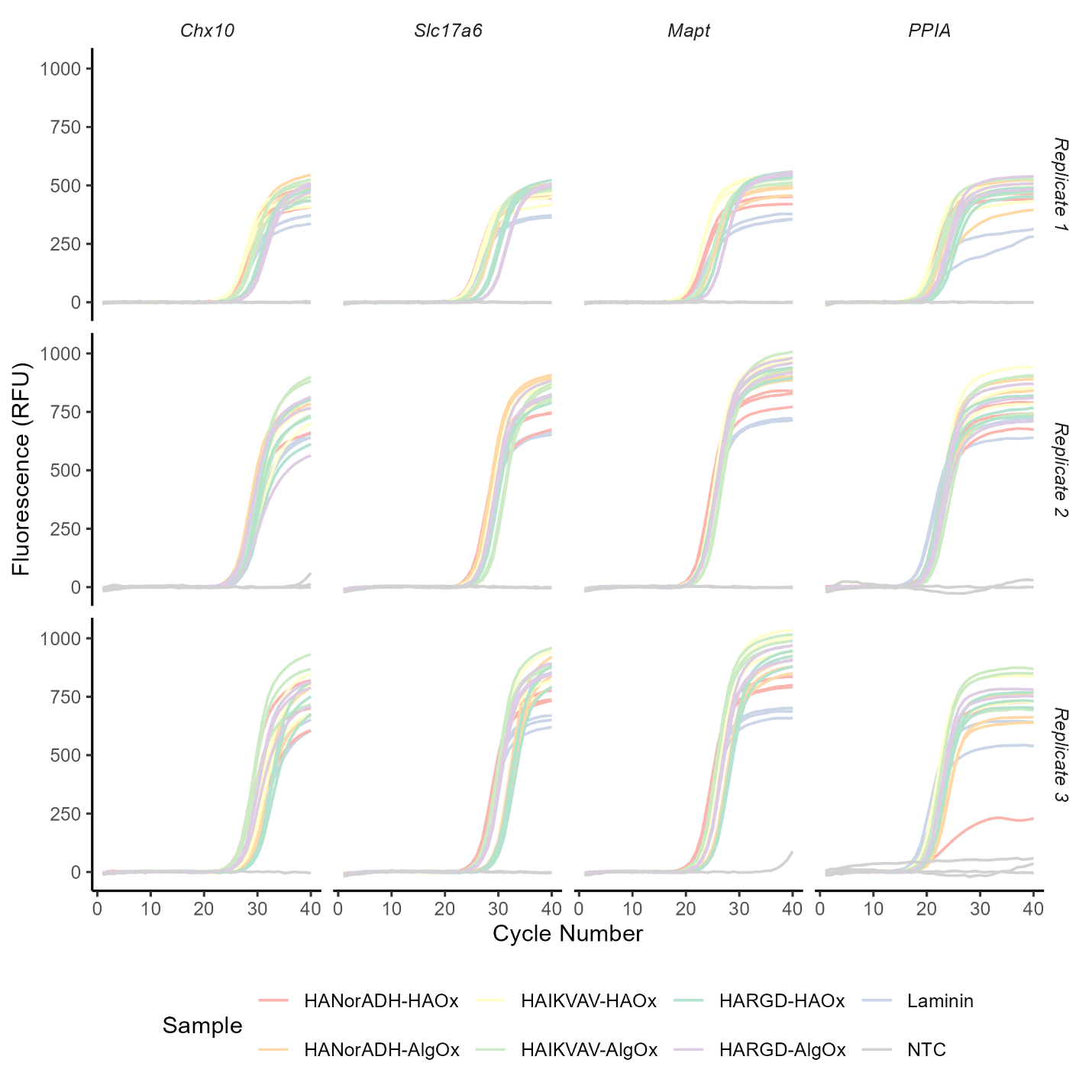
**Figure S4. Amplification profiles for qPCR.** Real-time amplification curves for target genes (*Chx10*, *Slc17a6*, *Mapt*, *Ppia*) exhibit characteristic sigmoidal profiles across hydrogel conditions and laminin controls. Flat profiles for non-template controls confirm the absence of genomic DNA contamination or primer-dimer artifacts. Columns correspond to a specific gene target, and rows represent independent biological replicates.

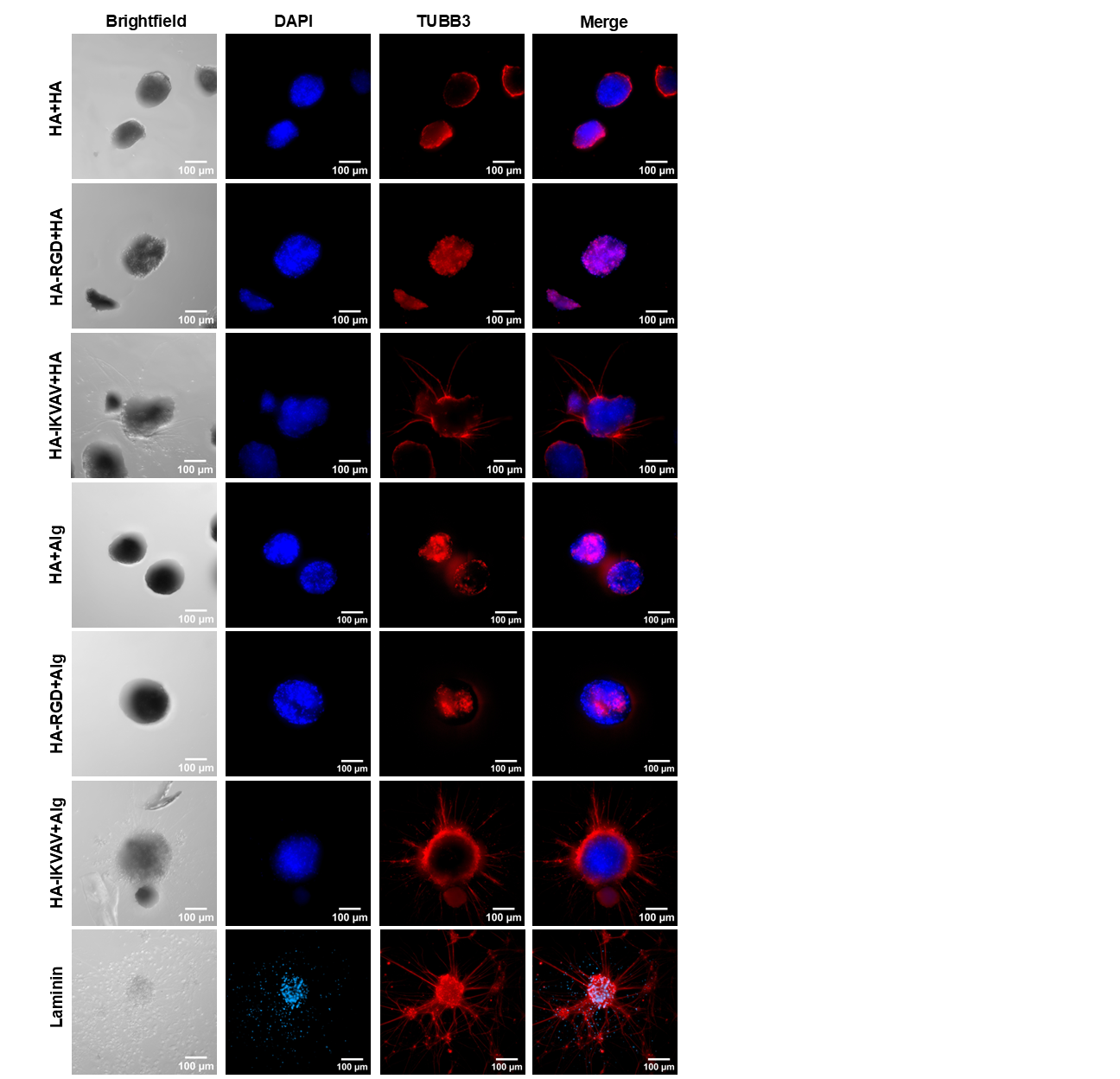

**Figure S5. Immunostaining of V2a interneurons**
Individual channels for immunostaining of V2a interneurons embedded in hydrogels and cultured on the 2D laminin control. Cells were immunostained for the neuronal microtubule marker βIII-Tubulin (TUBB3, red) and nuclei (DAPI, blue). Scale bars = 100 µm.
